# Deep learning based on stacked sparse autoencoder applied to viral genome classification of SARS-CoV-2 virus

**DOI:** 10.1101/2021.10.14.464414

**Authors:** Maria G. F. Coutinho, Gabriel B. M. Câmara, Raquel de M. Barbosa, Marcelo A. C. Fernandes

## Abstract

Since December 2019, the world has been intensely affected by the COVID-19 pandemic, caused by the SARS-CoV-2 virus, first identified in Wuhan, China. In the case of a novel virus identification, the early elucidation of taxonomic classification and origin of the virus genomic sequence is essential for strategic planning, containment, and treatments. Deep learning techniques have been successfully used in many viral classification problems associated with viral infections diagnosis, metagenomics, phylogenetic, and analysis. This work proposes to generate an efficient viral genome classifier for the SARS-CoV-2 virus using the deep neural network (DNN) based on stacked sparse autoencoder (SSAE) technique. We performed four different experiments to provide different levels of taxonomic classification of the SARS-CoV-2 virus. The confusion matrix presented the validation and test sets and the ROC curve for the validation set. In all experiments, the SSAE technique provided great performance results. In this work, we explored the utilization of image representations of the complete genome sequences as the SSAE input to provide a viral classification of the SARS-CoV-2. For that, a dataset based on *k*-mers image representation, with *k* = 6, was applied. The results indicated the applicability of using this deep learning technique in genome classification problems.

## 1. Introduction

Since the emergence of the SARS-CoV-2 virus at the end of 2019, many works are been developed aiming to provide more comprehension about this novel virus. In March 2020, the World Health Organization (WHO) raised the level of contamination to the COVID-19 pandemic, due to its geographical spread across several countries. On July 9, 2021, the disease had registered more than 185 million confirmed cases, and more than 4 million confirmed deaths. In the case of a novel virus identification, the early elucidation of taxonomic classification and origin of the virus genomic sequence is essential for strategic planning, containment, and treatments of the disease [1–3].

One of the fields of research in the bioinformatics area is the analysis of genomic sequences. In the last years, many strategies based on alignment-free methods have been explored as an alternative for the alignment-based methods, considering the limitations of the second approach. Alignment-based programs assume that homologous sequences comprise a series of linearly arranged and more or less conserved sequence stretches, which is not always the case in the real world [4].

Among the alignment-free methodologies, there are some models based on deep learning (DL) techniques, that can provide significant performance in applications of genome analysis [5–7]. Deep neural networks (DNN) can improve prediction accuracy by discovering relevant features of high complexity [7].

Figure 1 presents the genome analysis stages and how deep learning integrates this process. The genome analysis stages include the primary analysis, the secondary analysis, and the tertiary analysis. The primary and secondary analysis compose the genome sequencing. The primary analysis receives the biological sample and generates genomic data information, called “reads”, after the processing by the sequencer machine. Then, the secondary analysis processes the reads and produces the complete genome sequence. Lastly, the tertiary analysis provides the genome interpretation, which can be performed for many algorithms and techniques [8–10]. The deep learning techniques have been successful used for the tertiary analysis in many viral classification problems associated with the diagnosis of viral infections, metagenomics, pharmacogenomics, and others [11–15].

**Figure 1.**
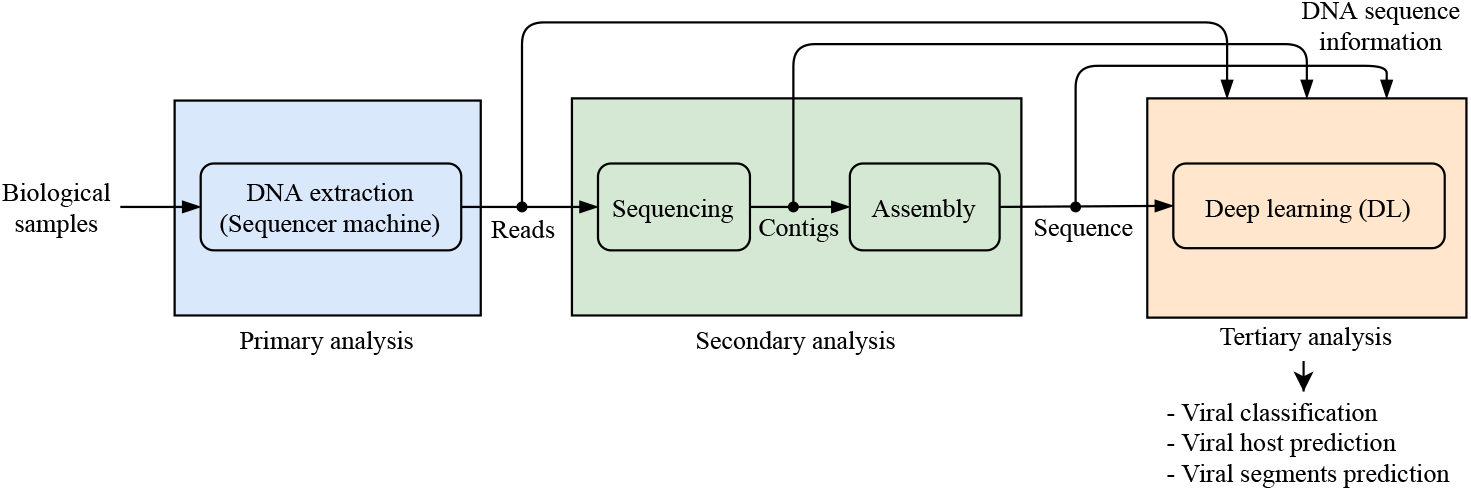
Genome analysis stages with deep learning.

Figure 2 shows the steps of the tertiary analysis using DL, that are the mapping and processing stages. The mapping stage receives the DNA sequence information, that can be the reads, contigs, or the whole genome sequence, and maps this data into a feature space. Various mapping strategies have been present in the works from the state of the art, such as one-hot encoding [13,16–18], number representation [11,12], digital signal processing [19], and other strategies, including multiple mapping strategies applied sequentially [20,21]. The processing stage consists of the utilization of a DNN to perform classification, prediction, and other assumptions about the genome information.

**Figure 2.**
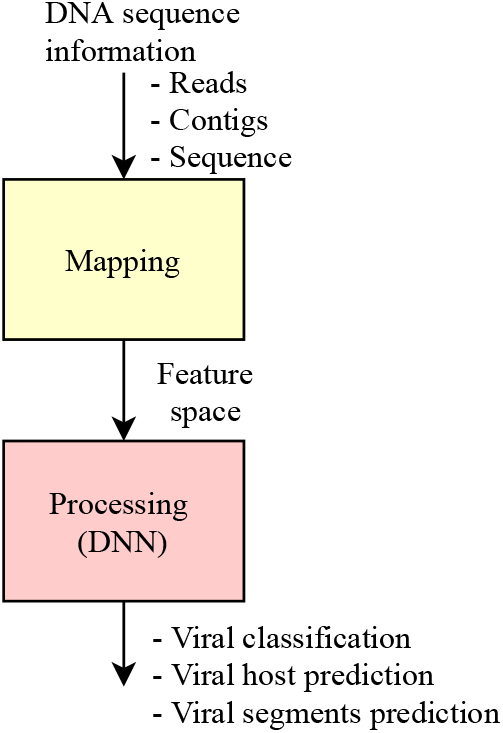
Stages of viral genome analysis using deep learning.

The mapping stage is crucial for the performance of the processing stage. The genome sequence length varies by the type of the virus. Since the DNN only receive a fixed-size input, some researchers have not been using the whole or long sequence length. Nevertheless, longer sequences contain more information and thus are more convenient to make predictions [17]. In this work, we will explore the utilization of the whole genome sequence mapped by image representation for the use as the DNN input in order to provide viral classification.

Recently works in literature have been applying deep learning as tertiary analysis such as viral prediction, viral host prediction, and viral segments prediction [11–19,22–30].

Tables 1 and 2 present some works from the state of the art that applied DNNs in order to analyse viral genome sequences. Table 1 details the focus of each work as the biology name, the group, the aim, indicates if the proposal was or was not applied for the COVID-19 and present the DNN used. The DNNs applied in those references are divide into 5 groups (CNN+FC, LSTM+FC, BLSTM+FC, BLSTM+CNN+FC, CNN+BLSTM+FC), as we show in the last column of Table 1. Table 2 shows the details about the input and the output of the DNN, besides the biology fields and the bioinformatics area.

**Table 1:**
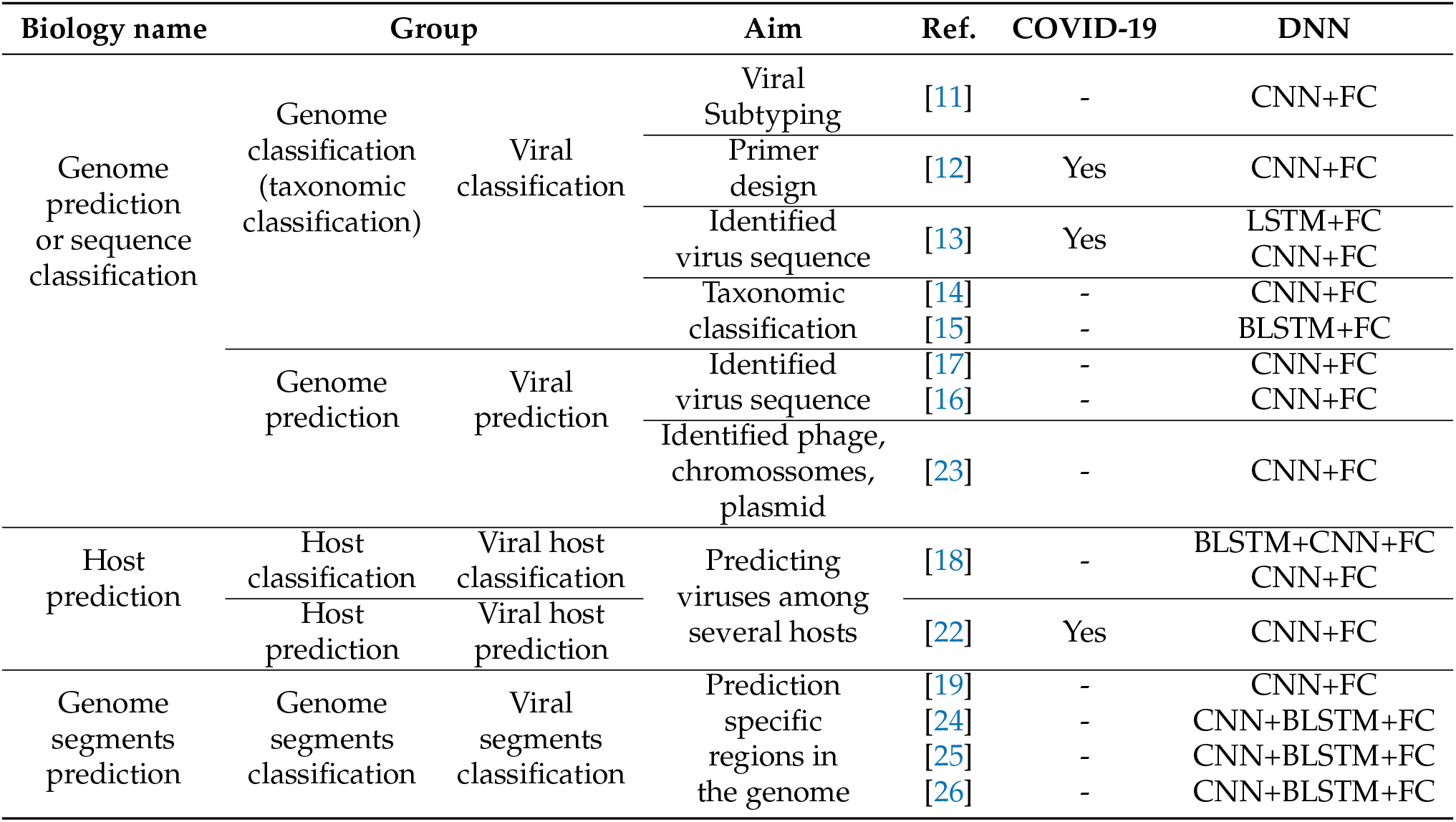
State of the art references - Part 1.

**Table 2:**
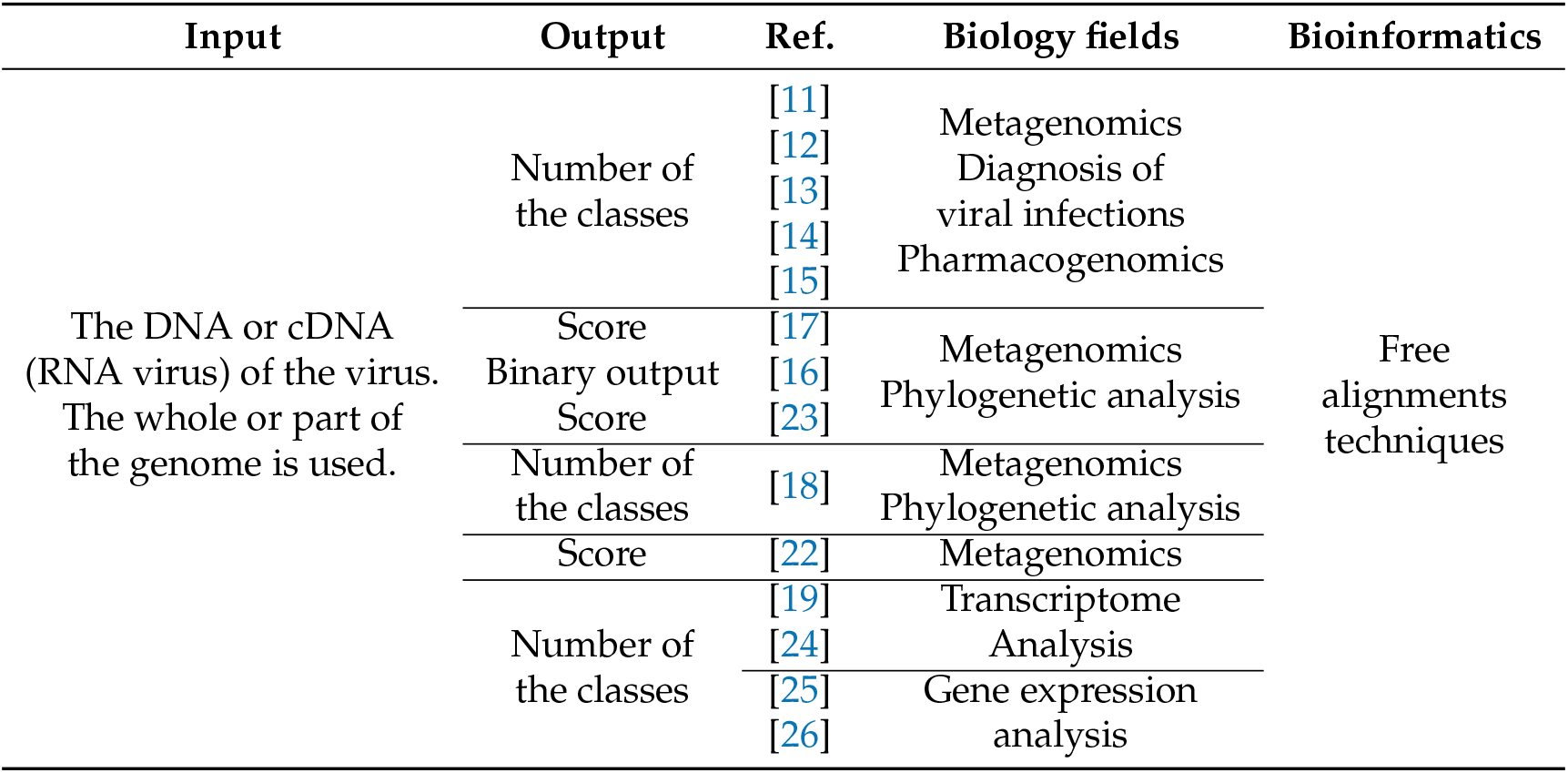
State of the art references - Part 2.

In the work presented in [11] was proposed a viral genome deep classifier (VGDC), the first viral genome subtyping based on deep learning techniques found in the literature. Their approach uses a Convolutional Neural Network (CNN) with 25 layers to classify several groups of viruses in subtypes. For the tests, were used five different datasets, each one containing genomes sequences of a specific type of virus. The whole virus genome sequence was used as the input to the network, where the corresponding ASCII code represented each nucleotide. The results indicated that the VGDC was able to achieve better results in comparison with previous works from the state of the art.

In [12] was proposed an approach to assist the tests in the detection of SARS-CoV-2, based on the use of DL techniques. For this, a CNN architecture with 4 layers was used to extract characteristics of the virus genomes, as well as to classify SARS-CoV-2 among Coronavirus type viruses. As presented in [11], the CNN received as input the whole virus genome sequences. The nucleotides were mapped in numerical values (C = 0.25, T = 0.50, G = 0.75, A = 1.0). Missing entries received a value of 0.0. The experiments showed that the CNN was able to correctly identify the sequences even in cases where the noise was added to the genome, reaching accuracies between 0.9674 (with noise) and 0.9875 (without noise). Through the results, the authors also identified a sequence as exclusive for the SARS-CoV-2 virus. They proposed the use of this sequence as a primer for PCR tests.

In [13], was proposed an approach to provide viral classification using the contigs (fragments of the genome sequence) and two different reverse-complement (RC) neural networks architectures: a RC-CNN and a RC-LSTM. These models were also applied to the SARS-CoV-2 virus.

In works presented in [14] and [15], a taxonomic classification for metagenomics applications is proposed. Both works used segments of genome (reads) with DL input (see Figure 1), and the output is the number of the classes. In [14], it was proposed two DL models, one to classify species, and another to classify genus. In [15], a hierarchical taxonomic classification for viral metagenomic data via DL, called CHEER, was proposed. Similar to the work proposed in [14], the CHEER framework classifies the genus, family, and genus.

Proposals presented in [16], [17] and [23] used the contigs with DL input for viral prediction, and classification. In [16], and [17] a DL virus identification framework was proposed and both cases try to recognize if the input is a virus or not.

In work from [16], called ViraMiner, was proposed and approach to detect the presence of viruses on raw metagenomic contigs from different human samples. They used a CNN architecture with two different convolutional branches (pattern and frequency branch) in order to extract relevant features. The outputs of these branches are concatenated and inserted into the fully connected (FC) layer. The ViraMiner output produces a single value that indicates the likelihood of the sequence belonging to the virus class.

In the proposal presented in [17], called DeepVirFinder, the output is a score between 0 and 1 for a binary classification between virus and prokaryote. They fragmented the genomes into non-overlapping sequences of different sizes (150, 300, 500, 1000, and 3000 bp). The sequences were mapped for the network input using the one-hot encoding method. Since they increase the length of the input, i.e. the sequence fragment, they achieve better performance results, which was measured by the area under the receiver operating characteristic curve (AUROC). The maximum AUROC achieved was 0.98 for the 3000 bp fragment.

The work presented in [23] identifies metagenomic fragments as phages, chromosomes or plasmids using the CNN technique. The experiments were performed using artificial contigs and real metagenomic data. The network output, provided by a softmax layer, consists of 3 scores that indicate the probability that each fragment belongs to a specific class.

In the works from [22] and [18] are present DL architectures for host prediction and classification. [22] used a CNN to provide host and infectivity prediction of SARS-CoV-2 virus. In [18] was proposed an approach to predict viral host from three different virus species (influenza A virus, rabies lyssavirus and rotavirus A) from the whole or only fractions of a given viral genome.

In the works from [19], [24], [25] and [26] were proposed methodologies to predict or classify specific regions in the genome sequence. [19] presented a methodology for the classification of three different functional genome types: coding regions, long noncoding regions, and pseudogenes in genomic data. They used a digital signal processing (DSP) methods, called Genomic signal processing (GSP), that converts the nucleotide sequence into a graphical representation of the information contained in the sequence. A CNN with 19 layers was used to perform the classification results.

The authors in [24] proposed a DL framework to identify similar patterns in DNA N6-methyladenine (6mA) sites prediction. This framework, called Deep6mA, is composed of a CNN to extract high-level features in the sequence and a Bi-directional LSTM (BLSTM) to learn dependence structure along the sequence, besides a fully connected layer that determines whether the site is a 6mA site.

In [25] was provided a method based on CNN and BLSTM for exploring the RNA recognition patterns of the CCCTC-binding factor (CTCF) and identify candidate IncR-NAs binding. The experiments conducted with two different datasets (human U2OS and mouse ESC) were able to predict CTCF-binding RNA sites from nucleotide sequences. Moreover, [26] propose a computational prediction approach for DNA–protein binding based on CNN and BLSTM.

We intend to provide viral classification using the whole genome sequences, as presented in [11] and [12]. However, in these works were used the length of the longest genome sequence of the dataset as the input of the DNN. So, it was necessary to add some padding for the missing entries. In this work, we will explore the utilization of *k*-mers image representation of the complete genome sequences as the DNN input, which will feasibly the use of genome sequences of any length and enable the use of smaller network inputs. The *k*-mers representation was used in many works that provide genome sequence classification, as presented in [31], which explores the spectral sequence representation based on *k*-mers occurrences. However, that work doesn’t explore the *k*-mers image representation.

We also explore the utilization of the stacked sparse autoencoder (SSAE) technique as an efficient viral genome classifier. The SSAE has been successfully applied in many biomedical works from the state of the art [6,32–34]. We performed some experiments to provide various levels of taxonomic classification of the SARS-CoV-2 virus, similar to the proposed experiments in [35], using the SSAE technique with a dataset of *k*-mers images representations, available on [36].

## 2. Materials and Methods

### 2.1. Dataset

For the experiments, we used a *k*-mers representation dataset of SARS-CoV-2 genome, available on [36]. This dataset is composed of 1557 virus instances of SARS-CoV-2, as also, a data stream of 11540 viruses from the Virus-Host DB dataset and the other three Riboviria viruses from NCBI (Betacoronavirus RaTG13, bat-SL-CoVZC45, and bat-SL-CoVZXC21). It also provides k-mers image representation of all data. The *k*-mers images were used to perform the experiments for this work.

Each *d*-th sequence, stored in dataset, is expressed by

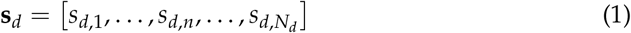

where *N*_*d*_ is the length of *d*-th sequence and *s*_*d,n*_ is the *n*-th nucleotide of the sequence. Each *n*-th *s*_*d,n*_ can be characterized as a symbol belonging to an alphabet of 4 possible symbols expressed by set {A, T, C, G} for DNA or by set {A, U, C, G} for RNA, that is,

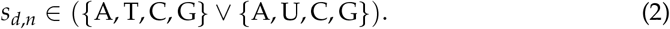

In *k*-mers representation, each *d*-th nucleotide sequence, **s**_*d*_, is grouped in *k*-mers sub-sequences [37,38] that can be expressed as

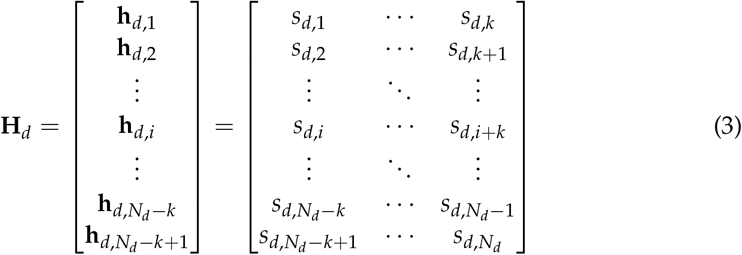

where the matrix **H**_*d*_ stores the *k*-mers associated with each *d*-th sequence **s**_*d*_. The *k*-mers representations are based in each *d*-th matrix **H**_*d*_ and the matrix **Γ**, call here as symbol matrix. The symbol matrix is expressed as

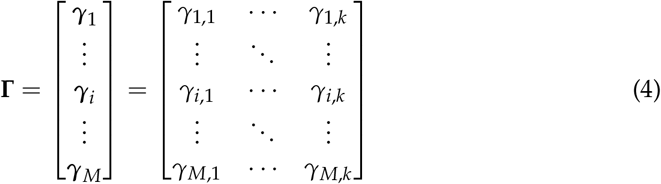

where each element *γ*_*i,j*_ ϵ ({A, T, C, G}∨{A, U, C, G}). The symbol matrix, **Γ**, stores all *M* possibilities of the *k*-mers, where

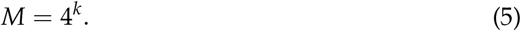

The *k*-mers count 1D representation can be expressed as

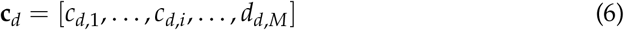

where

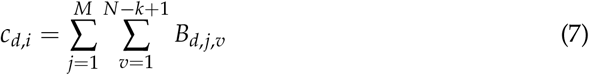

and

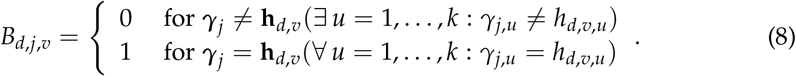

Table 3 shows a example of the *k*-mers count 1D representation values (with *k* = 2) for SARS-CoV-2 from China-Wuhan (ID: LR757995), USA-MA (ID: MT039888), Brazil (ID:MT126808), and Italy (ID: MT066156). The dataset provide in [36] has *k*-mers count 1D representation for *k* = 2, …, 6.

**Table 3:** Examples of *k*-mers count 1D representation values (with *k* = 2) for SARS-CoV-2.

| $k$-mers ($k = 2$) | China-Wuhan<br>(ID: LR757995) | USA-MA<br>(ID: MT039888) | Brazil<br>(ID: MT126808) | Italy<br>(ID: MT066156) |
| --- | --- | --- | --- | --- |
| AA | 2862 | 2859 | 2853 | 2847 |
| AC | 2022 | 2022 | 2022 | 2022 |
| AG | 1741 | 1741 | 1742 | 1742 |
| AT | 2306 | 2309 | 2309 | 2308 |
| CA | 2085 | 2082 | 2084 | 2082 |
| CC | 886 | 888 | 888 | 888 |
| CG | 439 | 439 | 440 | 439 |
| CT | 2080 | 2081 | 2080 | 2082 |
| GA | 1612 | 1612 | 1612 | 1611 |
| GC | 1167 | 1167 | 1169 | 1168 |
| GG | 1092 | 1093 | 1092 | 1092 |
| GT | 1990 | 1990 | 1988 | 1989 |
| TA | 2373 | 2378 | 2377 | 2378 |
| TC | 1415 | 1412 | 1413 | 1413 |
| TG | 2589 | 2589 | 2587 | 2587 |
| TT | 3212 | 3217 | 3219 | 3216 |

The *k*-mers count 2D representation for each *d*-th sequence, **s**_*d*_, is described by

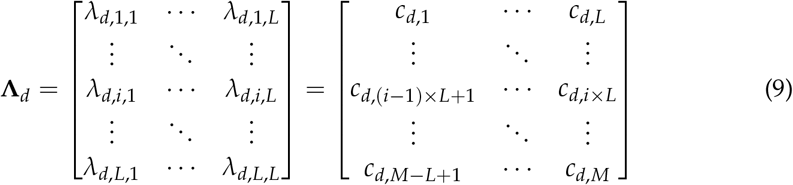

where

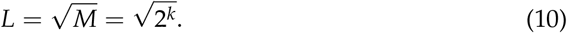

Finally, the *k*-mers image representation, for each *d*-th sequence, can be represented as

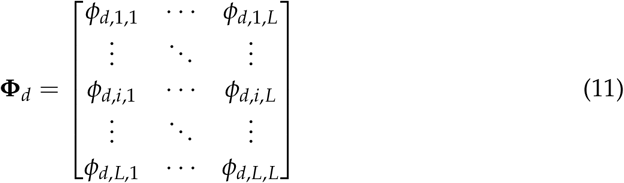

where *ϕ*_*d,i,j*_ represents each pixel associated with *d*-th image **Φ**_*d*_. Each pixel, *ϕ*_*d,i,j*_, is be expressed as

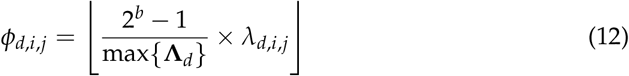

where max{·} is the maximum value in *d*-th matrix **Λ**_*d*_, ⌊·⌋is the greatest integer less than or equal, and *b* is number of bits associated with the image pixels. Figure 3 show the *k*-mers image representation, matrix **Φ**, (with *k* = 6 and *b* = 8) for Geminiviridae (ID: HE616777), Alphacoronavirus (ID: JQ410000), and SARS-CoV-2 (Betacoronavirus) from China-Wuhan (ID: LR757995) and Brazil (ID: MT126808).

**Figure 3.**
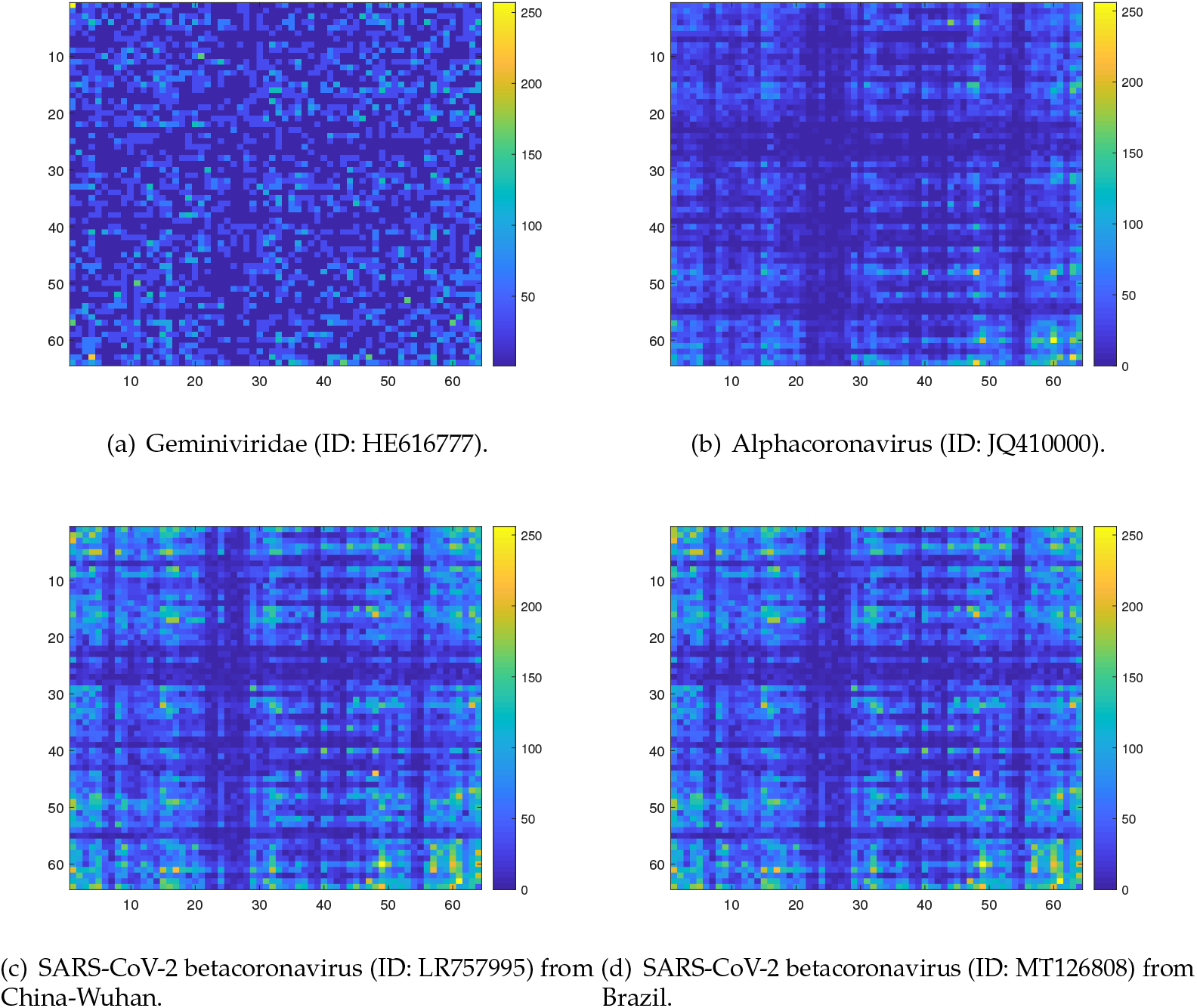
Examples of *k*-mers images representation with *k* = 6. Based on Equation 10, *L* = 64 and each image, matrix **Φ** (see Equation 11), is composed by 64 × 64 pixels with *b* = 8 (see Equation 12).

In this work, we used *k*-mers image representation with *k* = 6. In the work presented in [16], the 6-mers reached the best performance in comparisons with other values of *k* (3, 4, 5 and 7). The data of each experiment was partitioned using the holdout method, which splits the data into a training set and a validation set at random. We used the proportion of 80% for the training set and 20% for the validation set. Each class data was split respecting these percentages. The SARS-CoV-2 *k*-mers images were used only for the test set.

### 2.2. DNN Architecture

All experiments were performed using the SSAE technique. In these models each hidden layer is composed of an individually trained sparse autoencoder in an unsupervised way. A sparse autoencoder is an autoencoder whose training involves a sparse penalty, which functions as a regularizing term added to the loss function [39]. The autoencoder (AE) is a DL technique specialized in dimensionality reduction and feature extraction. The AE output can provide the reconstruction of the input information. These networks are composed of three layers: an input, a hidden and an output. The encoder is formed by the input and hidden layers, and the decoder is formed by the hidden and output layers [39]. For the output layer, we used a softmax layer, where the number of neurons consists of the number of classes of the experiment. Figure 4 illustrates the DL SSAE with *P* inputs, *K* hidden layers, and a output layer. Each *i*-th hidden layer has *Q*_*i*_ neurons and the output layer has *U* neurons. Functions *φ*(·) and *f* (·) are the action functions in each *p*-th neuron (in each *i*-th hidden layer) and each *u*-th neuron in output layer, respectively.

**Figure 4.**
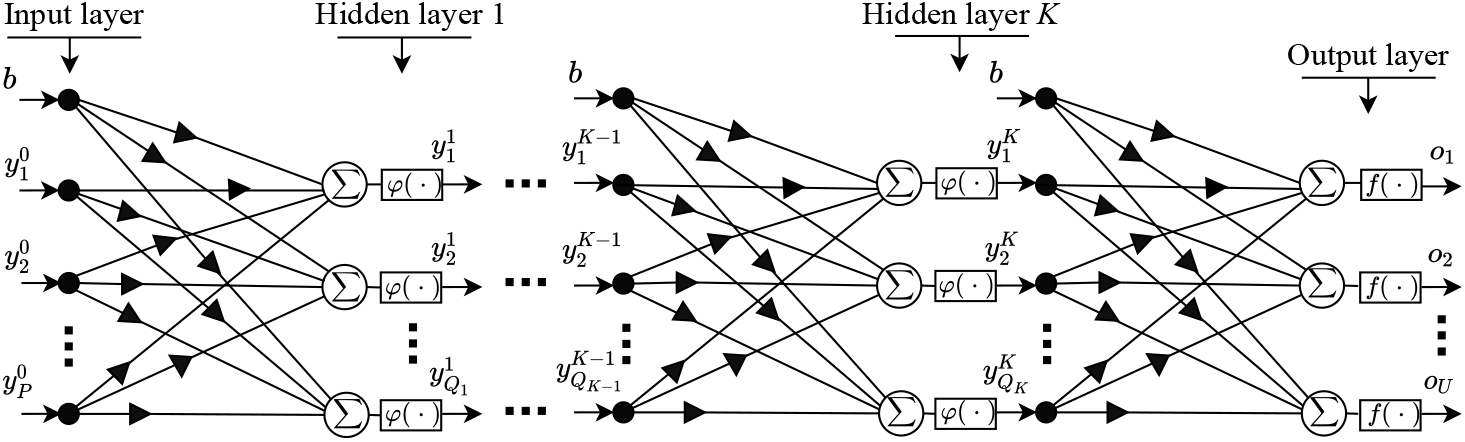
Deep learning stacked sparse autoencoder architecture (DL-SSAE).

For all experiments, the network architecture used three hidden layers (*K* = 3), containing 3000 neurons in the first hidden layer, *Q*_1_, 1000 in the second hidden layer, *Q*_2_, and 500 in the third hidden layer *Q*_3_. For input of the SSAE, it was used *k*-mers images, with *k* = 6, generating images, matrix **Φ**, with 64 × 64 pixels (based on Equation 10, 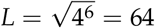). Each *d*-th image, **Φ***d*, associated with a *d*-th viral genome sequence is reshaped into a vector expressed by

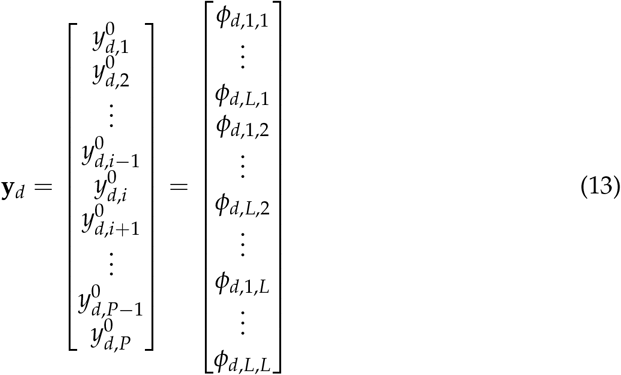

with *P* = 64 × 64 = 4096 values and applied to the SSAE. The number of neurons in output layer, *U*, is defined by the number of different viruses in a specific taxonomic level such as family, genus, realm and other. The output can be expressed by

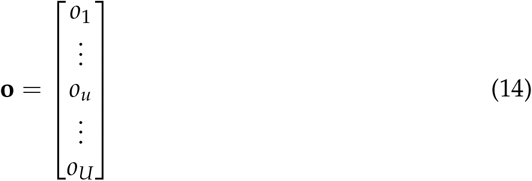

where each *u*-th output, *o*_*u*_, represents a specific virus in a taxonomic level classification and is defined by

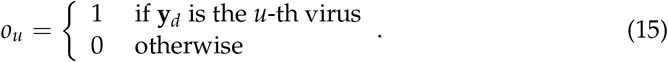

Figure 5 illustrates how the sequence information is passed through the DL-SSAE to perform the viral classification. The DL-SSAE input was normalized in the range of 0 to 1. First, the SSAE receives the training set as input to perform the training phase. Then, the validation set, which only contains samples that were not applied in the training phase, is used to identify the capacity of generalization of the DNN. After the network validation, the SSAE was applied for the test set, which only contains SARS-CoV-2 sequences. The SARS-CoV-2 *k*-mers images were not used for the training phase of the SSAE.

**Figure 5.**
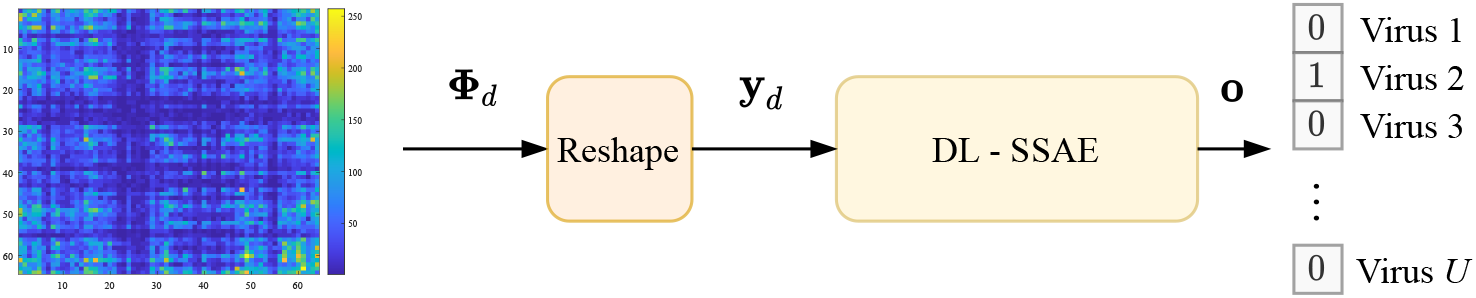
Viral classification process using *k*-mers images representation with the DL-SSAE.

The SSAE was implemented in the Matlab platform (License 596681) [40], adopting the deep learning toolbox. All network was trained with the Scaled Conjugate Gradient (SCG) algorithm. The loss function used for the training in each AE was the Mean Squared Error with L2 and Sparsity Regularizers, that can be expressed as

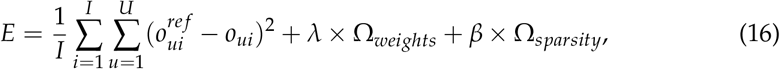

where *I* is the number of training examples, *U* is the number of classes, Ω_*weights*_ is the L2 regularization term, *λ* is the coefficient for the L2 regularization term, Ω_*sparsity*_ is the sparsity regularization term, and *β* is the coefficient for the sparsity regularization term.

The loss function applied for the softmax layer was the Cross-Entropy. In this work, after the training in each layer, the fine-tuning was performed, which retrained all the stacked network in a supervised way in order to improve the classification results. The fine-tuning process also used the Cross-Entropy as the loss function, as in the softmax layer.

## 3. Results and discussion

We performed four different experiments to provide different levels of taxonomic classification of the SARS-CoV-2 virus, similar to the experimental methodology present in [35]. The details about the data and the network architecture used in each experiment are shown in Table 5. The SSAE architecture was chosen by the observation of the MSE obtained with the reconstruction of the validation set in each AE. In order to validate the proposed idea of this work, the results are present by the confusion matrix for the validation and test sets. We also measured the performance of the viral classifier proposed with some popular classification metrics, as precision, recall, F1-score, and specificity. The precision value measure the percentages of all the examples predicted to belong to each class that are correctly classified, which corresponds to the positive predictive value. The recall, also called sensibility, corresponds to the percentages of all the examples belonging to each class that are correctly classified, which is the true positive rate. The F1-score can be interpreted as a weighted average of the precision and recall, and the specificity indicates the true negative rate. The column on the far right of each confusion matrix shows the percentages of precision per class, and the row at the bottom of each confusion matrix shows the percentages of recall per class. The cell in the bottom right of the plot of each confusion matrix shows the overall accuracy. Besides, for the validation set we also present the receiver operating characteristic (ROC) curve. The ROC curve measures the classification performance, that is the true positive rate and the false positive rate of each class, at various thresholds settings.

**Table 4:** Examples of *k*-mers count 2D representation values (with *k* = 2) for SARS-CoV-2.

| China-Wuhan (ID: LR757995) |  |  |  |  | USA-MA (ID: MT039888) |  |  |  |  |
| --- | --- | --- | --- | --- | --- | --- | --- | --- | --- |
| $\Lambda_{17} =$ | 2862 | 2022 | 1741 | 1741 | $\Lambda_{32} =$ | 2859 | 2022 | 1741 | 1741 |
|  | 2085 | 886 | 439 | 439 |  | 2082 | 888 | 439 | 439 |
|  | 1612 | 1167 | 1092 | 1092 |  | 1612 | 1167 | 1093 | 1093 |
|  | 2373 | 1415 | 2589 | 2589 |  | 2378 | 1412 | 2589 | 2589 |
| Brazil (ID: MT126808) |  |  |  |  | Italy (ID: MT066156) |  |  |  |  |
| $\Lambda_{52} =$ | 2853 | 2022 | 1742 | 1742 | $\Lambda_{79} =$ | 2853 | 2022 | 1742 | 1742 |
|  | 2084 | 888 | 440 | 440 |  | 2084 | 888 | 440 | 440 |
|  | 1612 | 1169 | 1092 | 1092 |  | 1612 | 1169 | 1092 | 1092 |
|  | 2377 | 1413 | 2587 | 2587 |  | 2377 | 1413 | 2587 | 2587 |

**Table 5:** Experiments data.

| Experiments | Classes | Number of sequences | SSAE architecture<br>$P - Q_1 - Q_2 - Q_3 - U$ |
| --- | --- | --- | --- |
| Experiment 1 | Adenoviridae | 195 | 4096 – 3000 – 1000 – 500 – 14 |
|  | Anelloviridae | 114 |  |
|  | Caudovirales | 500 |  |
|  | Circoviridae | 243 |  |
|  | Geminiviridae | 500 |  |
|  | Genomoviridae | 115 |  |
|  | Herpesvirales | 136 |  |
|  | Microviridae | 102 |  |
|  | Ortervirales | 214 |  |
|  | Papillomaviridae | 354 |  |
|  | Parvoviridae | 168 |  |
|  | Polyomaviridae | 142 |  |
|  | Riboviria | 500 |  |
|  | Tolecusatellitidae | 150 |  |
| Experiment 2 | Picornaviridae | 423 | 4096 – 3000 – 1000 – 500 – 8 |
|  | Caliciviridae | 392 |  |
|  | Coronaviridae | 206 |  |
|  | Potyviridae | 232 |  |
|  | Flaviviridae | 217 |  |
|  | Rhabdoviridae | 186 |  |
|  | Betaflexiviridae | 129 |  |
|  | Reoviridae | 111 |  |
| Experiment 3 | Alphacoronavirus | 52 | 4096 – 3000 – 1000 – 500 – 4 |
|  | Betacoronavirus | 123 |  |
|  | Deltacoronavirus | 20 |  |
|  | Gammacoronavirus | 9 |  |
| Experiment 4 | Embecovirus | 47 | 4096 – 3000 – 1000 – 500 – 4 |
|  | Merbecovirus | 17 |  |
|  | Nobecovirus | 9 |  |
|  | Sarbecovirus | 46 |  |

In Experiment 1, we intended to classify the viruses in 14 different classes, as presented in Table 5, which consists of 10 families (Adenoviridae, Anelloviridae, Circoviridae, Geminiviridae, Genomoviridae, Microviridae, Papillomaviridae, Parvoviridae, Polyomaviridae and Tolecusatellitidae), three orders (Caudovirales, Herpesvirales and Ortervirales) and Riboviria realm. The Riboviria class contains various families that belong to the realm Riboviria, including the Coronaviridae family. To ensure data balance, only the classes with at least 100 sequences from the original dataset were considered. For the classes with more than 500 sequences, only 500 sequences were selected at random, except for the Riboviria class, in which was prioritized the Coronaviridade family sequences, to guarantee the correct classification of the test data (SARS-CoV-2 sequences), which is the focus of this work. In this particular case, were selected all Coronaviridade family sequences available in the dataset (206 samples), and the other 294 sequences were select from the rest of the Riboviria data at random. After this balancing, Experiment 1 comprised 3433 samples of virus sequences.

The SSAE architecture used in Experiment 1 was the 4096 - 3000 - 1000 - 500 - 14 architecture. The three AEs were trained for 400 epochs. The softmax layer was trained for 3000 epochs or until reach the minimum gradient (< 1×10^−6^). Lastly, the fine-tuning was performed. For each experiment, the fine-tuning phase uses the same stopping condition as the softmax layer.

The confusion matrix and the ROC curve from the validation set of Experiment 1 are present in Figures 6 and 7, respectively. In Experiment 1, the classification accuracy from the validation set reached 92%. This result is promising, especially considering the challenges of the classification in high-level taxonomies because of the high diversity of the viruses sequences. It is essential to mention that the balancing process may have caused the classification more complicated because some crucial sequences may have been excluded from the dataset. However, this result can be improved in many ways that will be discussed following.

**Figure 6.**
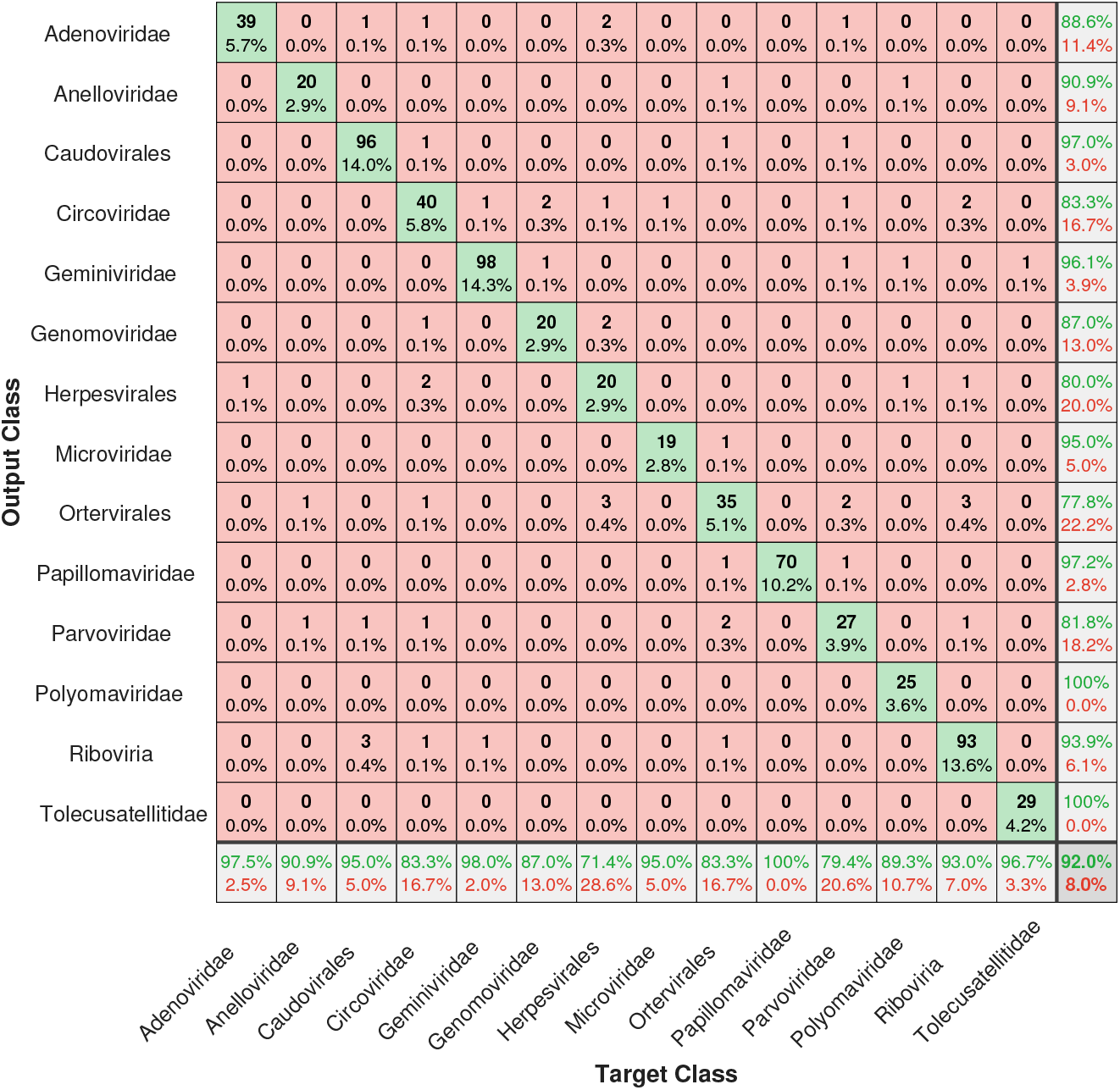
Confusion matrix of the validation set from the Experiment 1.

**Figure 7.**
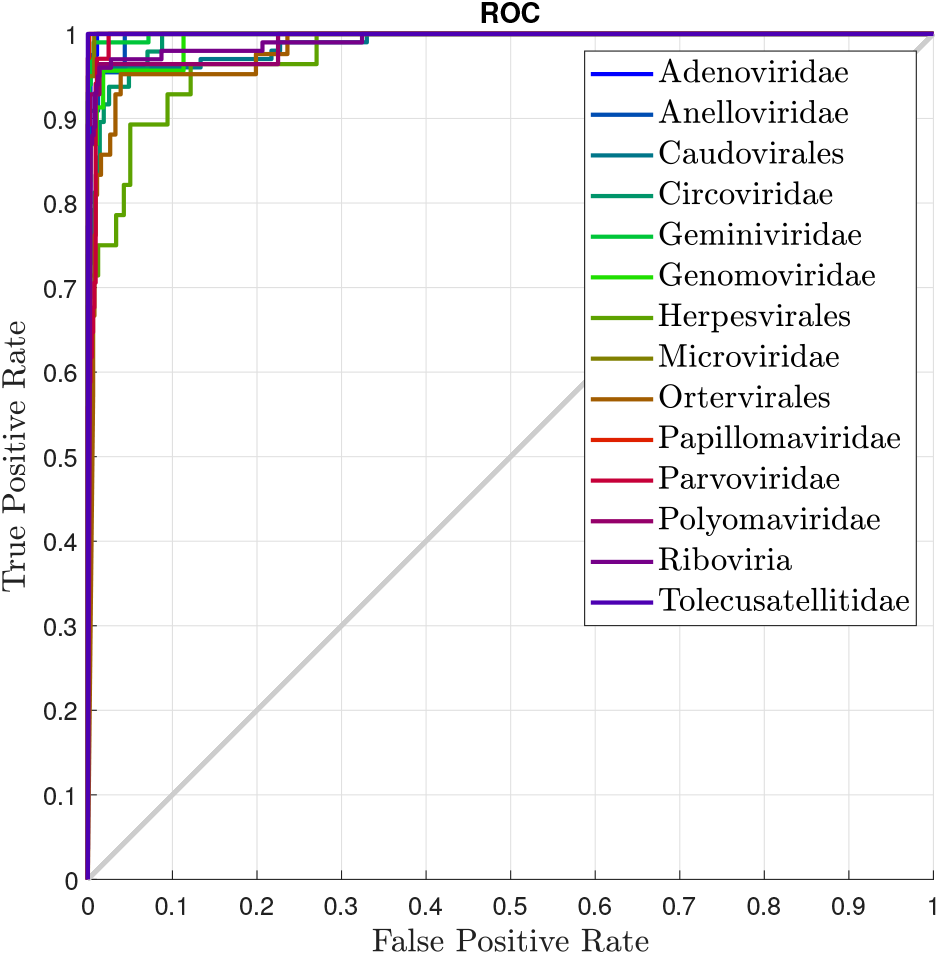
ROC curve of the validation set from the Experiment 1.

Regarded to the classification performance per class, the precision value presented in the last column shows that the worse result was obtained from an order class (71.4% from the Herpesvirales). Among the five worst classification results, two are from order classes (71.4% and 83.3% from Herpesvirales and Ortervirales, respectively). Since these classes can contain viruses from many different realms and families, they can difficult the training process. The Riboviria realm, which is the focus of this work, reached a classification accuracy of 93%. Analyse the results per classes can give more understanding about the dataset used and the implications of this dataset for the results, which is important to make decisions for the next experiments.

The confusion matrix from the test set of Experiment 1 is present in Figure 8. In the test phase of this experiment, all the 1557 sequences of SARS-CoV-2 was correctly classified as belonging to the Riboviria realm, so the classification accuracy reached 100%.

**Figure 8.**
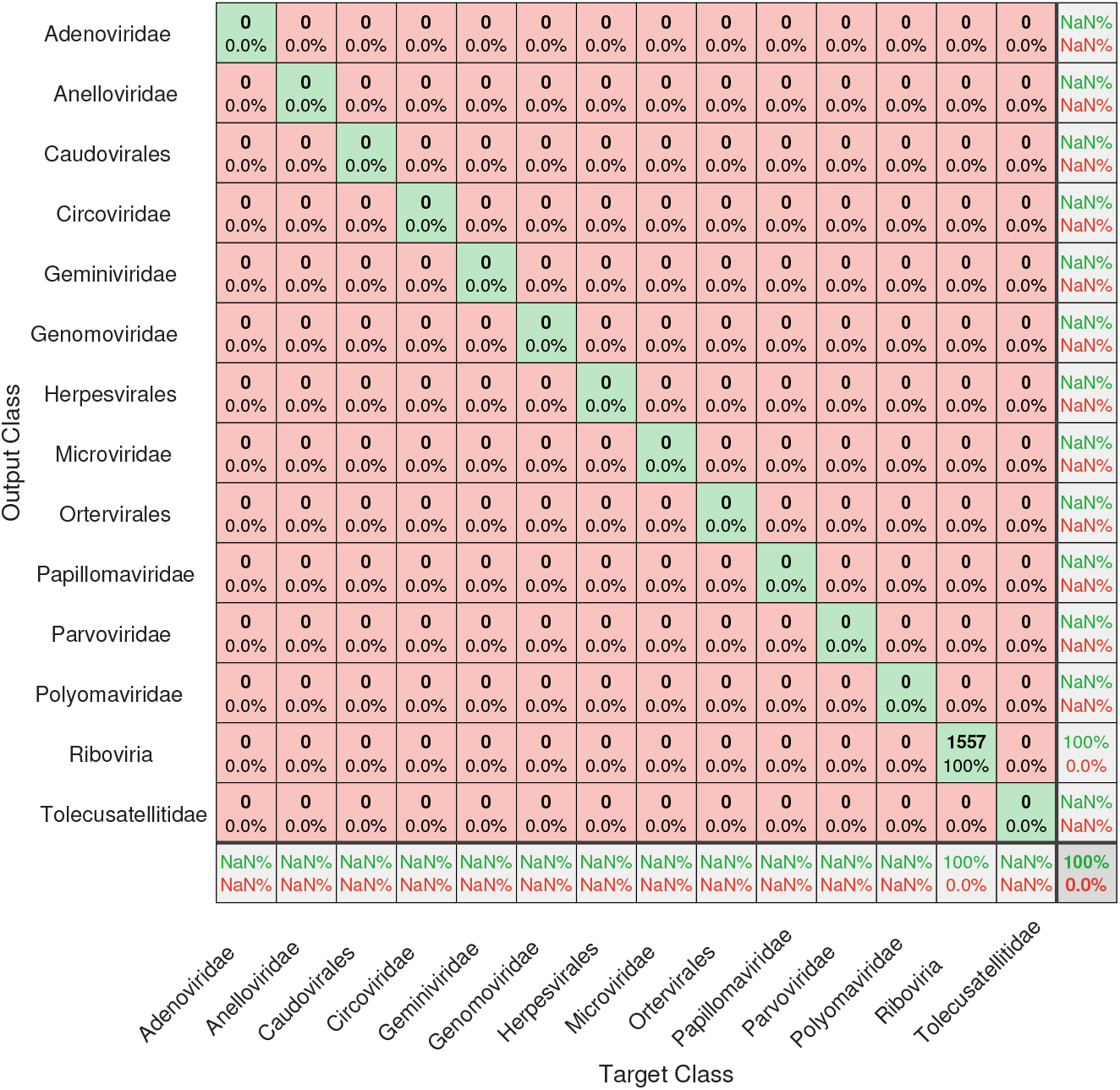
Confusion matrix of the test set from the Experiment 1.

Experiment 2 performs the classification of Riboviria families. As in Experiment 1, only classes with at least 100 sequences were considered. This experiment includes 1896 sequences separated into eight families (Picornaviridae, Caliciviridae, Coronaviridae, Potyviridae, Flaviviridae, Rhabdoviridae, Betaflexiviridae and Reoviridae). We used the 4096 - 3000 - 1000 - 500 - 8 SSAE architecture. The three AEs were trained for 400 epochs each and the softmax layer was trained for 1000 epochs or until reaching the minimum gradient, as well as the fine-tuning phase.

The confusion matrix and the ROC curve from the validation set of Experiment 2 are present in Figures 9 and 10, respectively. The classification accuracy from Experiment 2 reached 96.3%. From the 379 sequences applied in this validation, only 11 were not correctly classified. Besides, the SSAE classified all sequences that belong to the Coronaviridade family correctly. The ROC curve from Experiment 2 also provides excellent results.

**Figure 9.**
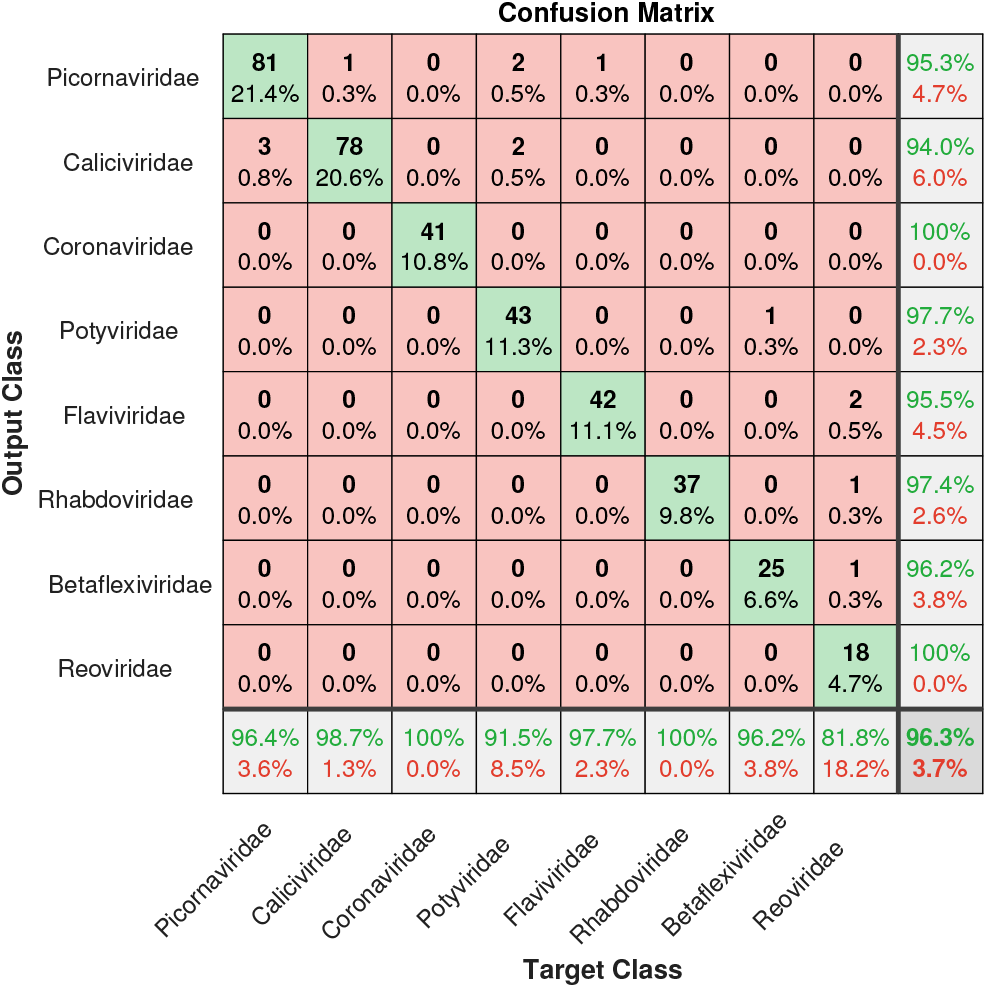
Confusion matrix of the validation set from the Experiment 2.

**Figure 10.**
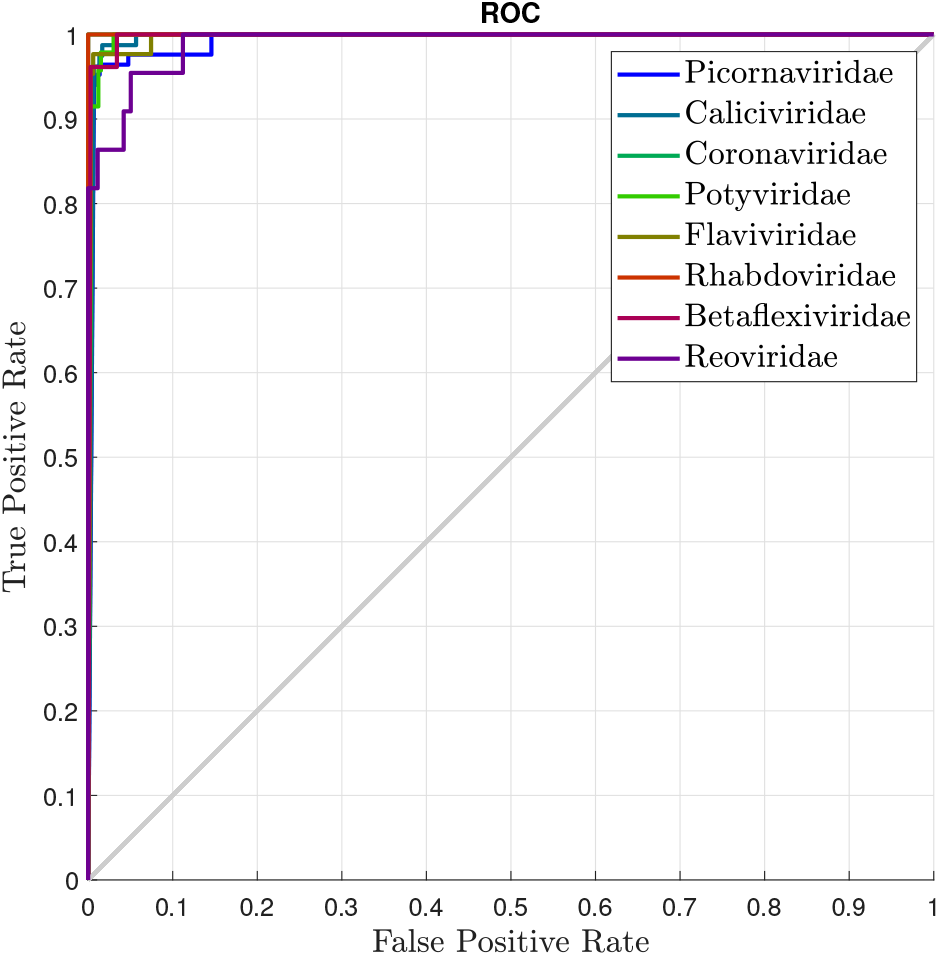
ROC curve of the validation set from the Experiment 2.

The confusion matrix from the test set of Experiment 2 is present in Figure 11. The SSAE achieve 100% of classification accuracy, i.e., all SARS-CoV-2 sequences applied in this experiment were perfectly classified as Coronaviridae family sequences.

**Figure 11.**
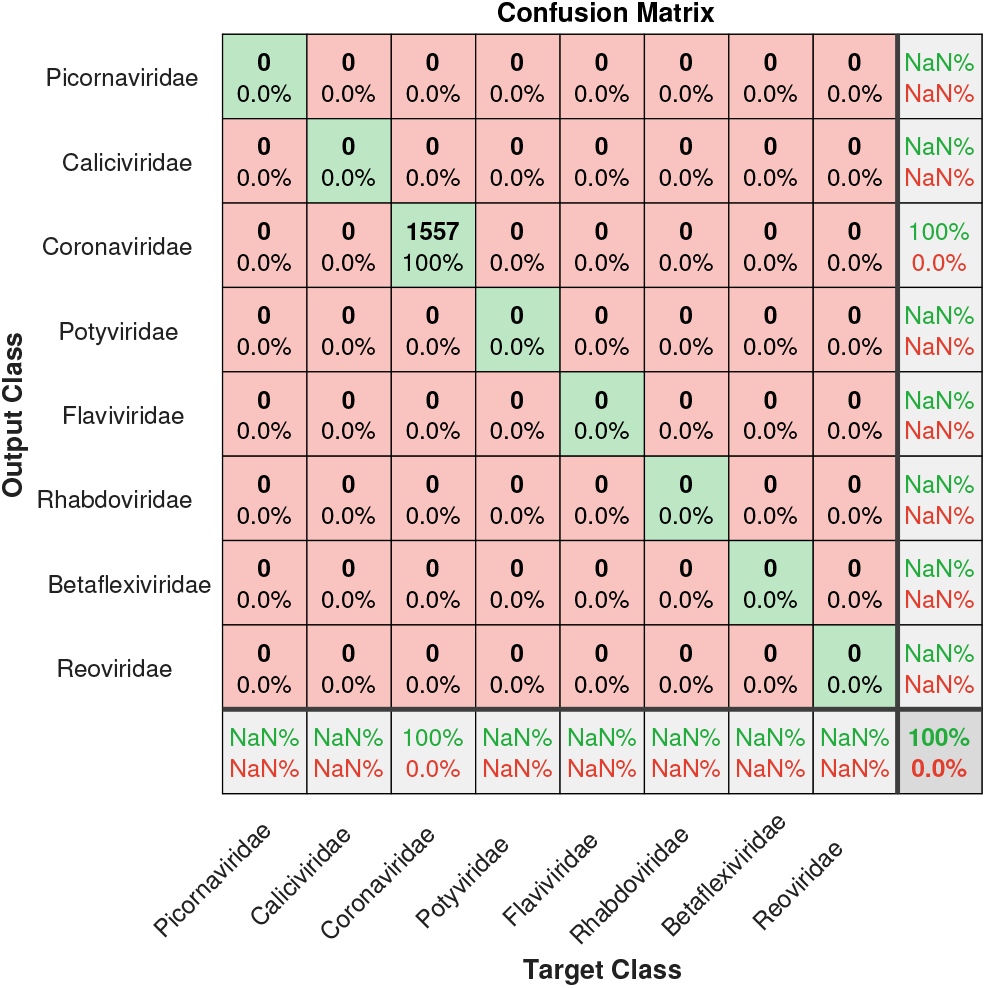
Confusion matrix of the test set from the Experiment 2.

In Experiment 3 we aim to provide the classification among the Coronaviridae genera. For this experiment, 204 sequences divided into four genera (Alphacoronavirus, Betacoronavirus, Deltacoronavirus and Gammacoronavirus) were used. The SSAE architecture used in this experiment was the 4096 - 3000 - 1000 - 500 - 4 architecture. The three AEs were trained for 400 epochs each, and the softmax layer was trained for 2000 epochs or until reaching the minimum gradient.

Figures 12 and 13 show the resulting confusion matrix and ROC curve from the Experiment 3, respectively. This experiment achieved 95% of classification accuracy of the validation set. The classification performance of the model obtained for the Betacoronavirus genus was 95.8%. Also, the ROC curve plotted for all classes of Experiment 3 provides satisfactory results.

**Figure 12.**
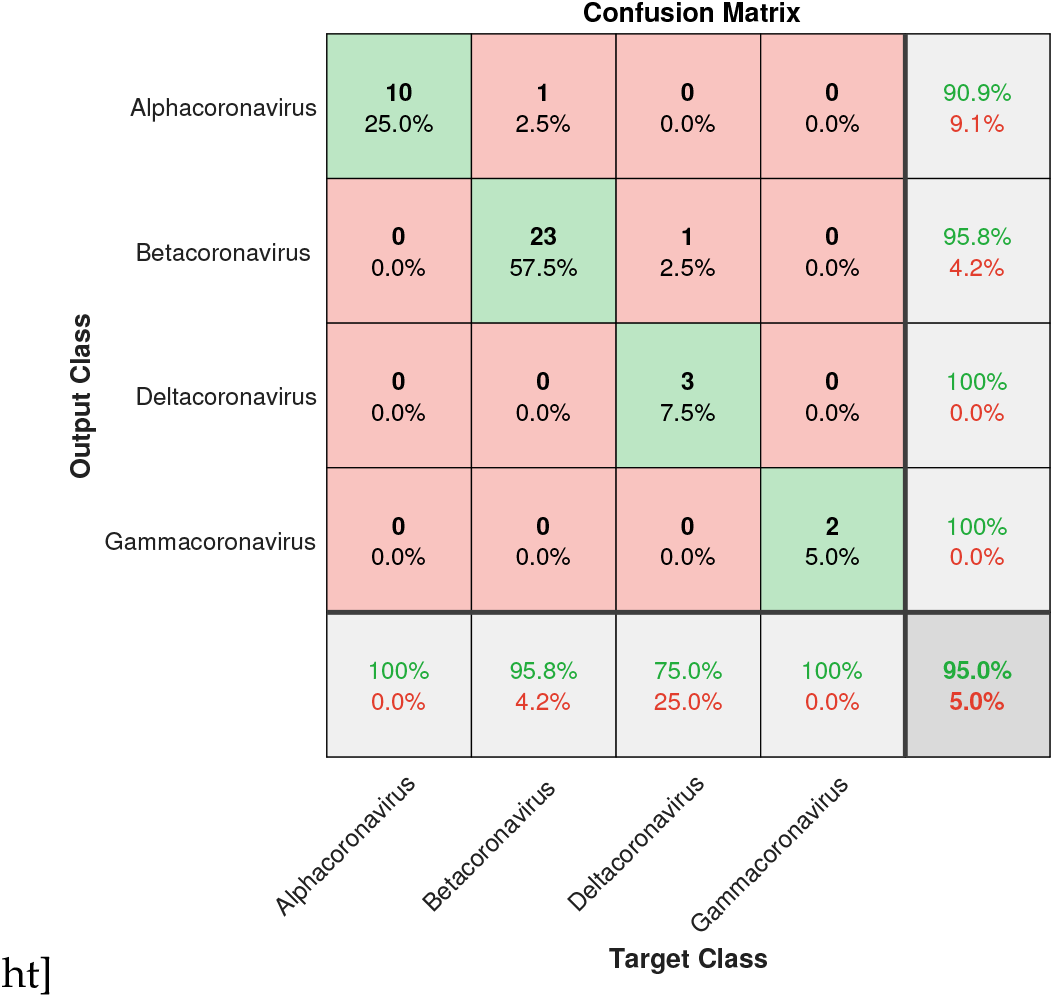
Confusion matrix of the validation set from the Experiment 3.

**Figure 13.**
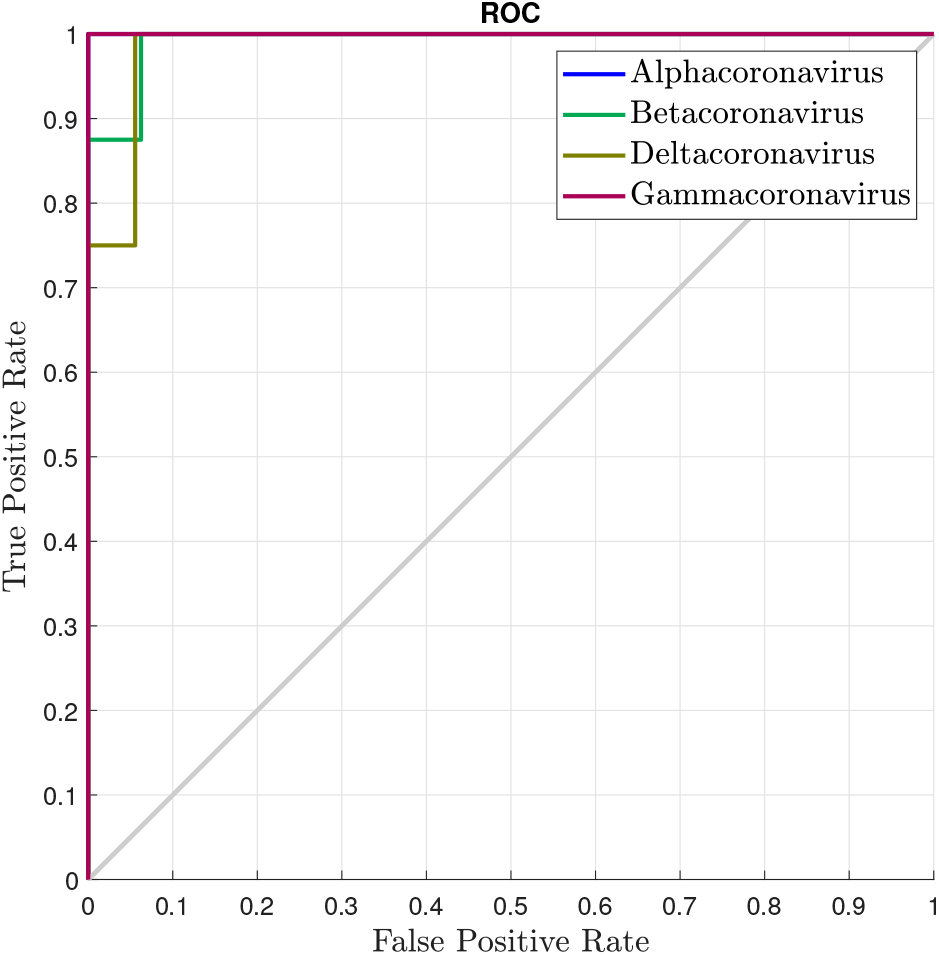
ROC curve of the validation set from the Experiment 3.

Regarding the test set of Experiment 3, the confusion matrix is present in Figure 14. The test phase of Experiment 3 achieved 98.9% of classification accuracy. In the validation phase of Experiment 3, the Betacoronavirus genus did not reach the highest performance, which probably explains these result in the test phase.

**Figure 14.**
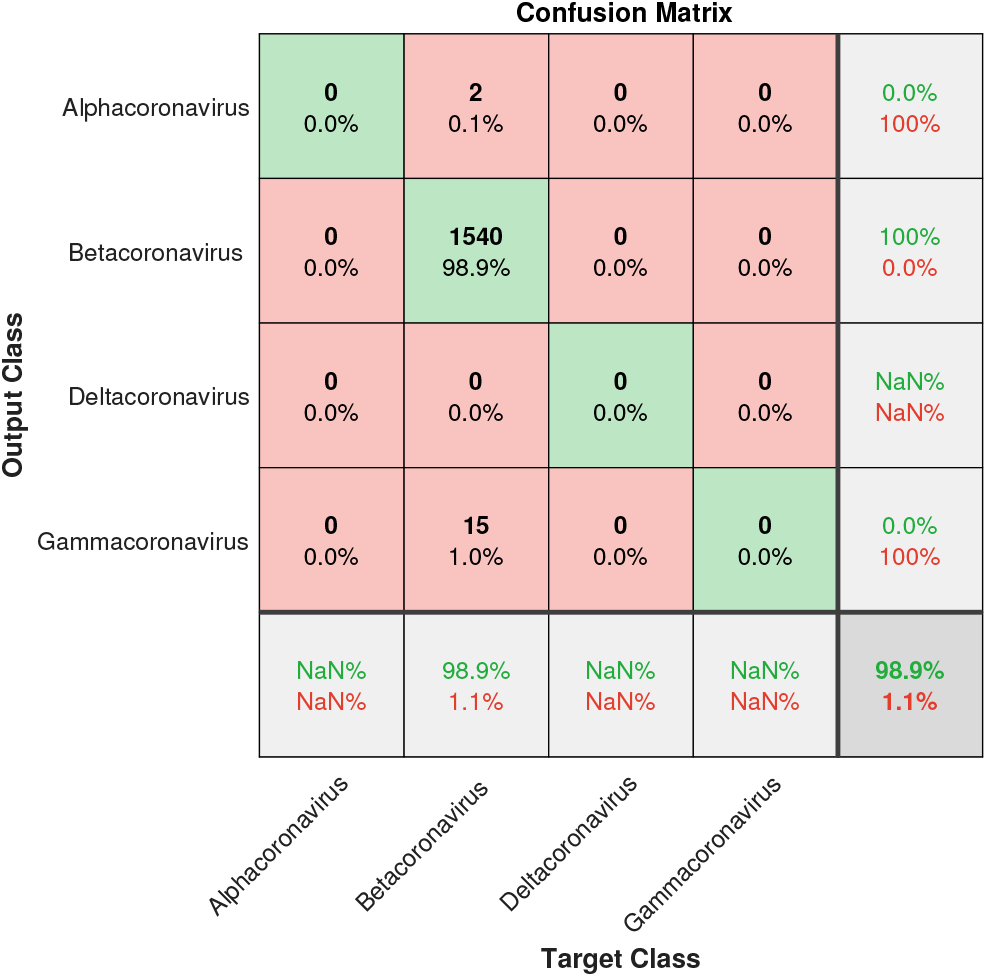
Confusion matrix of the test set from the Experiment 3.

In Experiment 4, we provide the Betacoronaviridae subgenera classification. This test includes 119 genome sequences divided into four classes (Embecovirus, Marbecovirus, Nobecovirus and Sarbecovirus). The SSAE architecture was the same as the architecture used in Experiment 3 (4096 - 3000 - 1000 - 500 - 4), as well as the training parameters.

The confusion matrix and the ROC curve from the validation set of Experiment 4 are present in Figures 15 and 16, respectively. In this experiment, the SSAE achieved the highest classification accuracy (100%), which is reaffirmed for the ROC curve plot.

**Figure 15.**
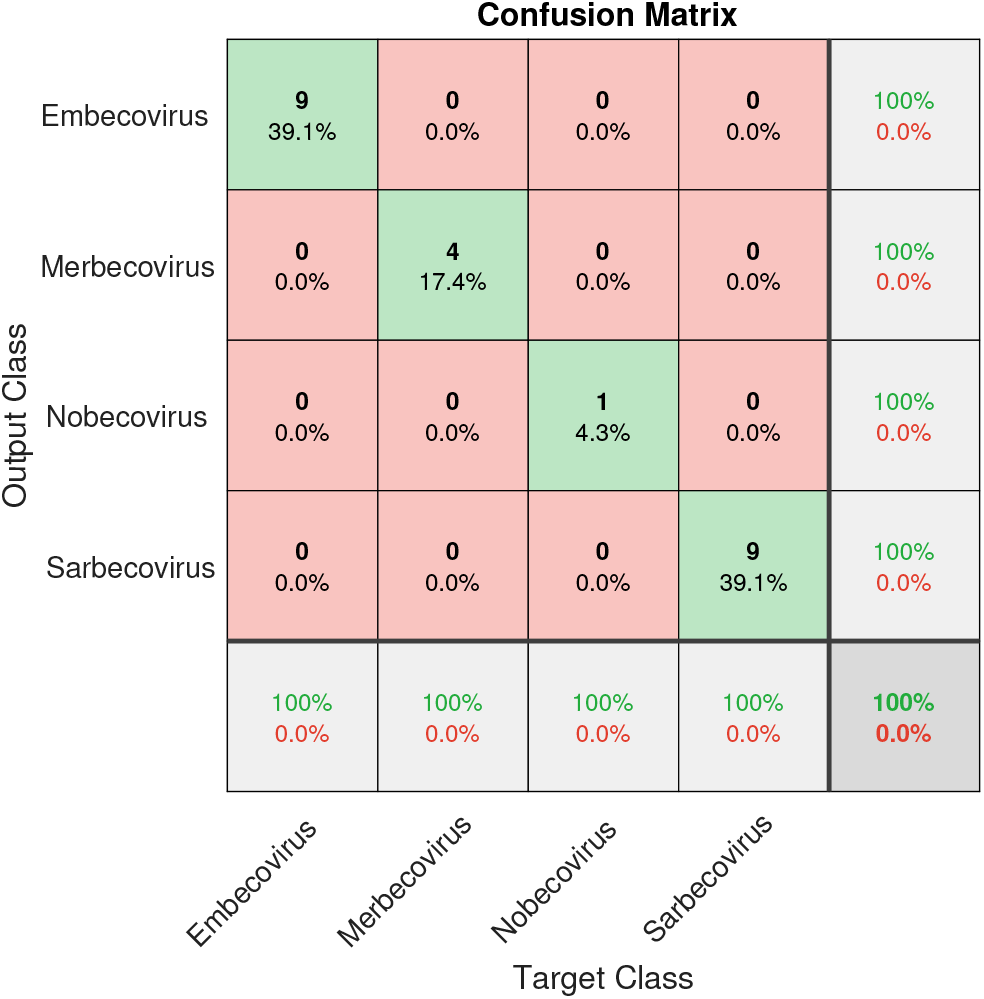
Confusion matrix of the validation set from the Experiment 4.

**Figure 16.**
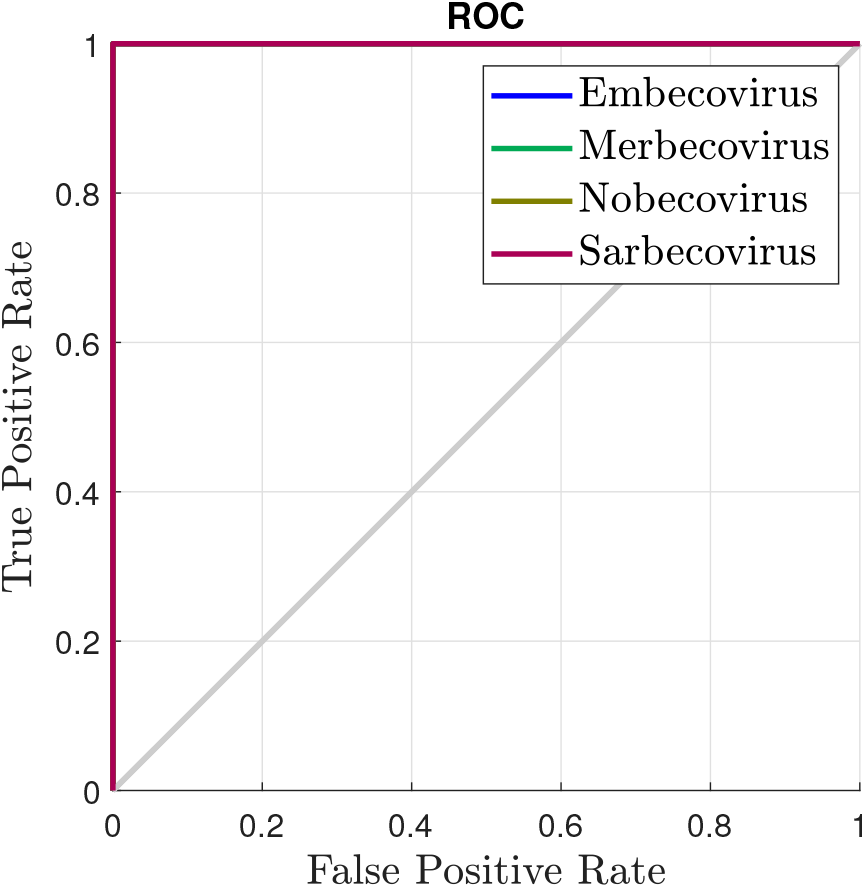
ROC curve of the validation set from the Experiment 4.

**Figure 17.**
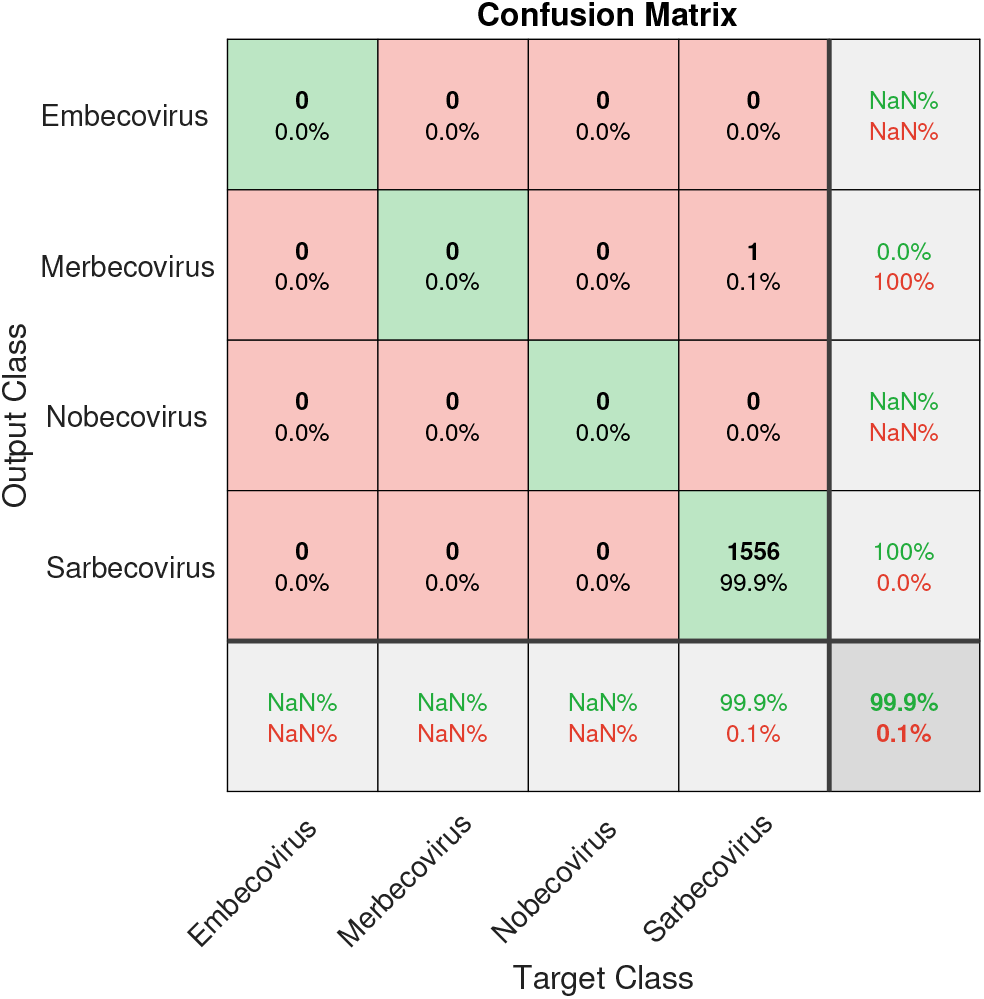
Confusion matrix of the test set from the Experiment 4.

Figure 15 exposes the confusion matrix from the test set of Experiment 4. In this case, the SSAE achieved 99.9% of classification accuracy, that is equivalent to only one sequence wrong classified.

Table 6 presents the results regarding some popular classification performance metrics obtained from the validation set. The first column of the table indicates the experiment proposed. The second column shows the overall accuracy for each experiment. The precision, recall, F1-score, and specificity are present in the others columns, which were obtained by the average of the values obtained for each class.

**Table 6:**
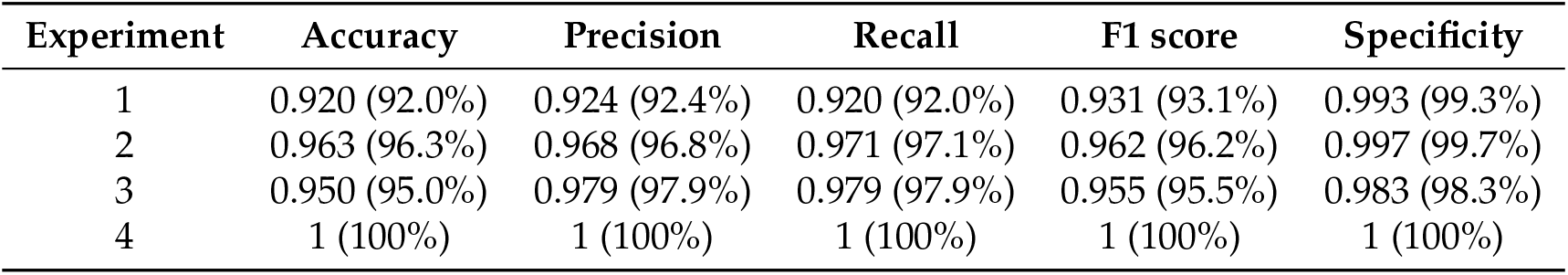
Classification performance metrics results obtained from the validation set.

All the metrics presented in Table 6 indicate that the viral classifier proposed performs great for all experiments. The highest performance was obtained for the Experiment 4. Besides, Experiments 2 and 3, reached values more than 0.95 for all the metrics evaluated. The classification performance slightly decreased in the Experiment 1, which is acceptable because of the high diversity of the viruses sequences applied. However, considering all the experiments, the specificity (true negative rate) reached values between 0.983 and 1.

Table 7 presents the results regarding some popular classification performance metrics obtained from the test set. The first column of the table indicates the experiment proposed. The second column shows the overall accuracy for each experiment. And the last column shows the recall, or true positive rate, which were obtained only for the class that corresponds to the SARS-CoV-2 samples. The other metrics (precision, F1-score, and specificity) are not presented because in the tests we do not have false positives samples.

**Table 7:**
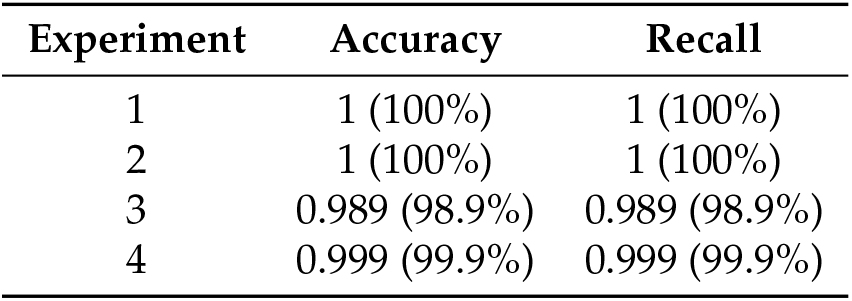
Classification performance metrics results obtained from the test set.

When the SARS-CoV-2 samples were applied, all the experiments perform excellently. The accuracy reached values between 98.9% and 100%, as well as the recall (true positive rate). The results presented in Table 7 are very significant since the classification of the SARS-CoV-2 virus was the main objective of this study.

In all experiments, the SSAE technique provided great performance results, especially for the test set. However, some strategies can be applied in future experiments to improve classification accuracy results. One of them consists in the use of the *k*-fold cross-validation scheme. Besides, we also intend to study data balancing alternatives, based on the analysis of the results presented here.

## 4. Conclusions

This work presented an alignment-free methodology, based on the stacked sparse autoencoder technique, in order to classify genome sequences of the SARS-CoV-2 virus in various levels of taxonomy (realm, family, genus and subgenus). We explored the utilization of *k*-mers image representation of the whole genome sequence, which feasibility the use of genome sequences of any length and enable the use of smaller network inputs. The results were presented by the confusion matrix for the validation and test sets, and the ROC curve for the validation set. All experiments provided great performance results, reaching accuracies between 98.9% and 100% for the test set. These results indicated the applicability of using the stacked sparse autoencoder technique in genome classification problems.

## Author Contributions

All the authors have contributed in various degrees to ensure the quality of this work. (e.g., Maria G. F. Coutinho, Gabriel B. M. Câmara, Raquel de M. Barbosa and Marcelo A. C. Fernandes conceived the idea and experiments; Maria G. F. Coutinho, Gabriel B. M. Câmara, Raquel de M. Barbosa and Marcelo A. C. Fernandes designed and performed the experiments; Maria G. F. Coutinho, Gabriel B. M. Câmara, Raquel de M. Barbosa and Marcelo A. C. Fernandes analyzed the data; Maria G. F. Coutinho, Gabriel B. M. Câmara, Raquel de M. Barbosa and Marcelo A. C. Fernandes wrote the paper. Marcelo A. C. Fernandes coordinated the project.). All authors have read and agreed to the published version of the manuscript.

## Acknowledgments

The authors wish to acknowledge the financial support of the Coordenação de Aperfeiçoamento de Pessoal de Nível Superior (CAPES) for their financial support.

